# Global Geographic and Temporal Analysis of SARS-CoV-2 Haplotypes Normalized by COVID-19 Cases during the Pandemic

**DOI:** 10.1101/2020.07.12.199414

**Authors:** Santiago Justo Arevalo, Daniela Zapata Sifuentes, Cesar Huallpa Robles, Gianfranco Landa Bianchi, Adriana Castillo Chavez, Romina Garavito-Salini Casas, Guillermo Uceda-Campos, Roberto Pineda Chavarria

## Abstract

Since the identification of SARS-CoV-2, a large number of genomes have been sequenced with unprecedented speed around the world. This marks a unique opportunity to analyze virus spreading and evolution in a worldwide context. Currently, there is not a useful haplotype description to help to track important and globally scattered mutations. Also, differences in the number of sequenced genomes between countries and/or months make it difficult to identify the emergence of haplotypes in regions where few genomes are sequenced but a large number of cases are reported. We propose an approach based on the normalization by COVID-19 cases of relative frequencies of mutations using all the available data to identify major haplotypes. Furthermore, we can use a similar normalization approach to tracking the temporal and geographic distribution of haplotypes in the world. Using 171 461 genomes, we identify five major haplotypes (OTUs) based on nine high-frequency mutations. OTU_3 characterized by mutations R203K and G204R is currently the most frequent haplotype circulating in four of the six continents analyzed. On the other hand, during almost all months analyzed, OTU_5 characterized by the mutation T85I in nsp2 is the most frequent in North America. Recently (since September), OTU_2 has been established as the most frequent in Europe. OTU_1, the ancestor haplotype is near to extinction showed by its low number of isolations since May. Also, we analyzed whether age, gender, or patient status is more related to a specific OTU. We did not find OTU’s preference for any age group, gender, or patient status. Finally, we discuss structural and functional hypotheses in the most frequently identified mutations, none of those mutations show a clear effect on the transmissibility or pathogenicity.

## INTRODUCTION

COVID-19 was declared a pandemic by the World Health Organization on March 11^th^, 2020 (Cuccinota and Vanelli, 2020), with around 71 million cases and 1.6 million deaths around the world (December 14^th^) (WHO, 2020), quickly becoming the most important health concern in the world. Several efforts to produce vaccines, drugs, and diagnostic tests to help in the fight against SARS-CoV-2 are being mounted in a large number of laboratories all around the world.

Since the publication on January 24^th^ of the first complete genome sequence of SARS-CoV-2 from China (Zhu et al. 2020), thousands of genomes have been sequenced in a great number of countries on all 5 continents and were made available in several databases. This marks a milestone in scientific history and gives us an unprecedented opportunity to study how a specific virus evolves in a worldwide context. As of November 30^th^, 2020, the GISAID database (Shu et al. 2017) contained 171 461 genomes with at least 29 000 sequenced bases.

Several analyses have been performed to identify SARS-CoV-2 variants around the world, most of them on a particular group of genomes using limited datasets (For example, Saha et al. 2020, Maitra et al. 2020, Castillo et al. 2020, Franco-Muñoz et al. 2020). In March 2020 two major lineages were proposed based on position 8782 and 28144 using a data set of 103 genomes (Tang et al. 2020) which was followed by a particularly interesting proposal that identified the same major lineages (named A and B) and other sublineages (Rambaut et al. 2020).

To complement these current classification systems, we consider that haplotypes description and nomenclature could help to better track important mutations that are currently circulating in the world. Identification of SARS-CoV-2 haplotypes aids in understanding the evolution of the virus and may improve our efforts to control the disease.

To perform a reasonable analysis of the worldwide temporal and geographical distribution of SARS-CoV-2 haplotypes, we need to take into account the differences in the number of sequenced genomes in months and countries or continents. Thus, we first used a data set of 171 461 complete genomes to estimate the worldwide relative frequency of nucleotides in each SARS-CoV-2 genomic position and found nine mutations with respect to the reference genome EPI_ISL_402125 with normalized relative frequencies (NRFp) representing to be present in more than 9 500 000 COVID-19 cases. After that, using a total of 109 953 complete genomes without ambiguous nucleotides from GISAID we performed a phylogenetic analysis and correlated the major branches with SARS-CoV-2 variants which can be classified into five haplotypes or Operational Taxonomic Units (OTUs) based on the distribution of the nine identified nucleotide positions in our NRFp analysis. After that, we analyzed the geographical and temporal worldwide distribution of OTUs normalized by the number of COVID-19 cases. Also, we attempt to correlate these OTUs with patient status, age, and gender information. Finally, we discuss the current hypothesis of the most frequent mutations on protein structure and function. All this information will be continuously updated in our publicly available web-page (http://sarscov2haplofinder.urp.edu.pe/).

## RESULTS AND DISCUSSION

### Mutations frequency analysis

The GISAID database contains 171 461 genomes with at least 29 000 sequenced bases; from these, 109 953 genomes do not present ambiguities (as of November 30th). With an alignment of the 171 461 genomes, we performed a normalized relative frequency analysis of each nucleotide in each genomic position (NRFp) (see material and methods for details). This normalization was performed to detect relevant mutations that could appear in regions where few genomes were sequenced (Fig. S1 shows that no correlation exists between the number of cases and the number of sequenced genomes). Using this NRFp analysis, we identified nine positions estimated to be in more than 9 500 000 COVID-19 cases (more than 0.18 NRFp) (Fig. 1.A and S2.A) plus many other mutations with NRFp between 0.00-0.18 (Fig. S2.B and S2.C).

**Figure 1.**
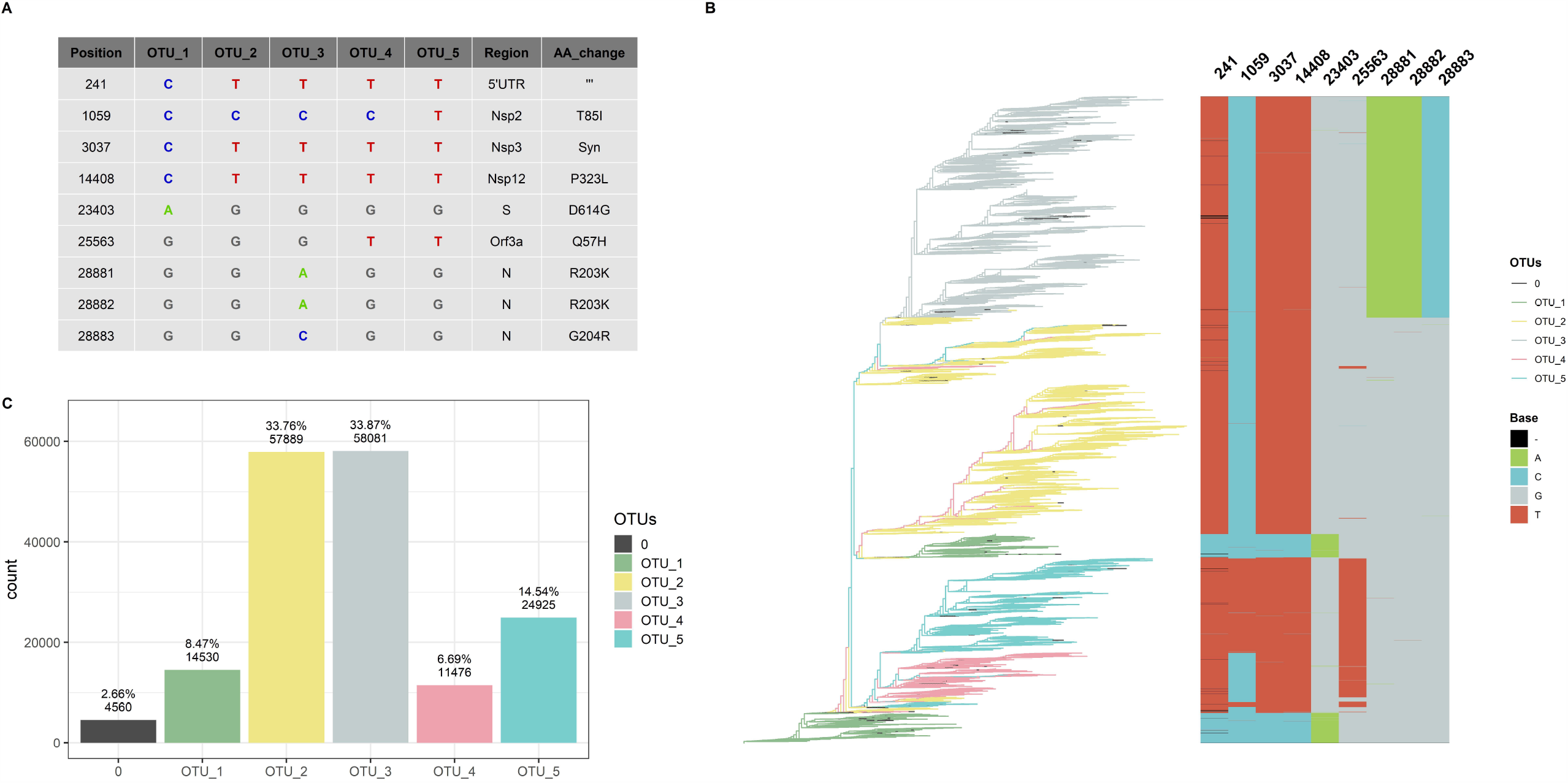
Five haplotypes (or OTUs) based in nine positions can classify 97 % of the genomes. A) Table showing haplotype of each OTU, regions, and aminoacids changes caused by these mutations. B) Rooted tree of 109 953 SARS-CoV-2 complete and non-ambiguous genomes associated with an alignment of nine genomic positions (241, 1059, 3037, 14408, 23403, 25563, 28881, 28882, 28883) showing a good correlation between haplotypes (OTUs) based in these nine positions. Tips of the tree where colored based in the OTU. C) Bar diagram showing OTUs distribution of the genomes (0 correspond to unclassified genomes).

The nine most frequent mutations (NRFp greater than 0.18) comprise seven non-synonymous mutations, one synonymous mutation, and one mutation in the 5’-UTR region of the SARS-CoV-2 genome (Fig. 1.A). The three consecutive mutations G28881A, G28882A, and G28883C falls at the 5’ ends of the forward primer of “China-CDC-N” (Table. S1). Because these three mutations are at the 5’ ends, it is unlikely that those mutations greatly affect amplification efficiency. The other six mutations do not fall within regions used by qRT-PCR diagnostic kits (Table. S1). All these nine mutations have been already identified in other studies (Korber et al. 2020, Kern et al. 2020, Pachetti et al. 2020, Yin et al. 2020), although with different frequencies mainly due to the absence of normalization.

### OTUs identification

After NRFp analysis, we estimated a maximum likelihood tree using the whole-genome alignment of the 109 953 complete genomes without ambiguities. Then, we associated the branches of the tree with an alignment of the nine positions (241, 1059, 3037, 14408, 23403, 25563, 28881, 28882, 28883). We noted that combinations of those nine positions represent five well-defined groups in the tree (Fig. 1.B). Using these combinations, we defined 5 haplotypes that allow us to classify more than 97 % of the analyzed genomes (Fig. 1.C), a great part of the remaining not classified genomes are due to the absence of sequencing corresponding to position 241. We named these haplotypes Operational Taxonomic Units (OTUs).

OTU_1 was considered the ancestor haplotype due to its identity with the first isolated genomes (EPI_ISL_402125 and EPI_ISL_406801) with characteristic C241, C3037, C14408, and A23403. This OTU_1 comprised genomes with T or C in position 8782 and C or T in 28144. In other analyses, these mutations divide SARS-CoV-2 strains into two lineages. For instance; at the beginning of the pandemic, Tang et al. (2020) showed linkage disequilibrium between those positions and named them as S and L lineages. Rambaut et al. (2020) used these positions to discriminate between their proposed major lineages A and B. Those mutations did not reach the estimated number of 9 500 000 COVID-19 cases, indicating that a small number of these genomes emerged during the pandemic in comparison with other variations.

A SARS-CoV-2 isolated on January 25th in Australia is at present the first belonging to OTU_2 (Fig. S3). Showing simultaneously four mutations different to OTU_1 (C241T, C3037T, C14408T, and A23403G), OTU_2 is the first group containing the D614G and the P323L mutations in the spike and nsp12 protein, respectively. Korber et al. (2020) analyzed the temporal and geographic distribution of this mutation separating SARS-CoV-2 into two groups, those with D614 and those with G614. Tomaszewski et al. (2020) analyzed the entropy of variation of these two mutations (D614G and P323L) until May. Apparently, OTU_2 is the ancestor of two other OTUs (OTU_3 and OTU_4), as shown in the maximum likelihood tree (Fig. 1.B). OTU_2 is divided into two major branches, one that originates OTU_3 and another more recent branch characteristic from Europe (see below, worldwide geographical distribution of OTUs).

On February 16th in the United Kingdom, a SARS-CoV-2 with three adjacent mutations (G28881A, G28882A, and G28883C) (Fig. S3) in N protein was isolated. These three mutations (together with those that characterized OTU_2) define OTU_3. The maximum likelihood tree shows that OTU_4 comes from OTU_2. OTU_4 does not present mutations in N protein; instead, it presents a variation in Orf3a (G25563T). Finally, OTU_5 presents all the mutations of OTU_4 plus one nsp2 mutation (C1059T).

These nine mutations have been separately described in other reports but, to our knowledge, they have not yet used been used together to classify SARS-CoV-2 haplotypes during the pandemic. The change of relative frequencies of those mutations analyzed individually showed that just in few cases, mutations that define haplotypes described here appear independently (Fig. S4). For example, the four mutations that define OTU_2 (C241T, C30307T, C14408T and A23403G) rarely had been described separately and similarly with mutations that characterize OTU_3 (G28881A, G28882A, G28883C) (Fig. S4). Thus, in this case analysis of haplotypes will be identical results that if we analyzed those mutations independently.

The fact that we were able to classify more than 97 % of the complete genomes data set (Fig. 1.C) shows that, at least to the present date, this classification system covers almost all the currently known genomic information around the world. Also, most of the unclassified tips appear within a clade allowing us to easily establish their phylogenetic relationships to a haplotype. Thus, at the moment this system can be of practical use to analyze the geographical and temporal distribution of haplotypes during these eleven months of 2020. For convenience we presented table S2 that contains the relation between our identified OTUs and their relationships with pangolin lineages (Rambaut et al. 2020) and GISAID clades (Shu et al. 2017).

### Worldwide geographic distribution of OTUs

Using our OTUs classification, we analyzed the worldwide geographic distribution during eleven months of 2020. We began by plotting continental information in the ML tree of the unambiguous complete genomes (Fig. 2.A) and observed some interesting patterns. For instance, all continents contain all OTUs; also, is relatively clear that most isolates belonging to OTU_5 come from North America (Fig. 2.A). Furthermore, the biggest branch of OTU_2 is almost exclusively filled by genomes from Europe, is interesting to note that this branch also contains genomes isolated in the last months analyzed showing its relatively recent appearance (see below, the worldwide temporal distribution of OTUs). However, this approach does not allow us to evaluate continents with less sequenced genomes (Fig. S5.A), such as South America, Oceania, and Africa. Also, it is possible that fine differences can be found in the frequency of one OTU concerning another in each continent. These differences are not observed at this level of analysis.

**Figure 2.**
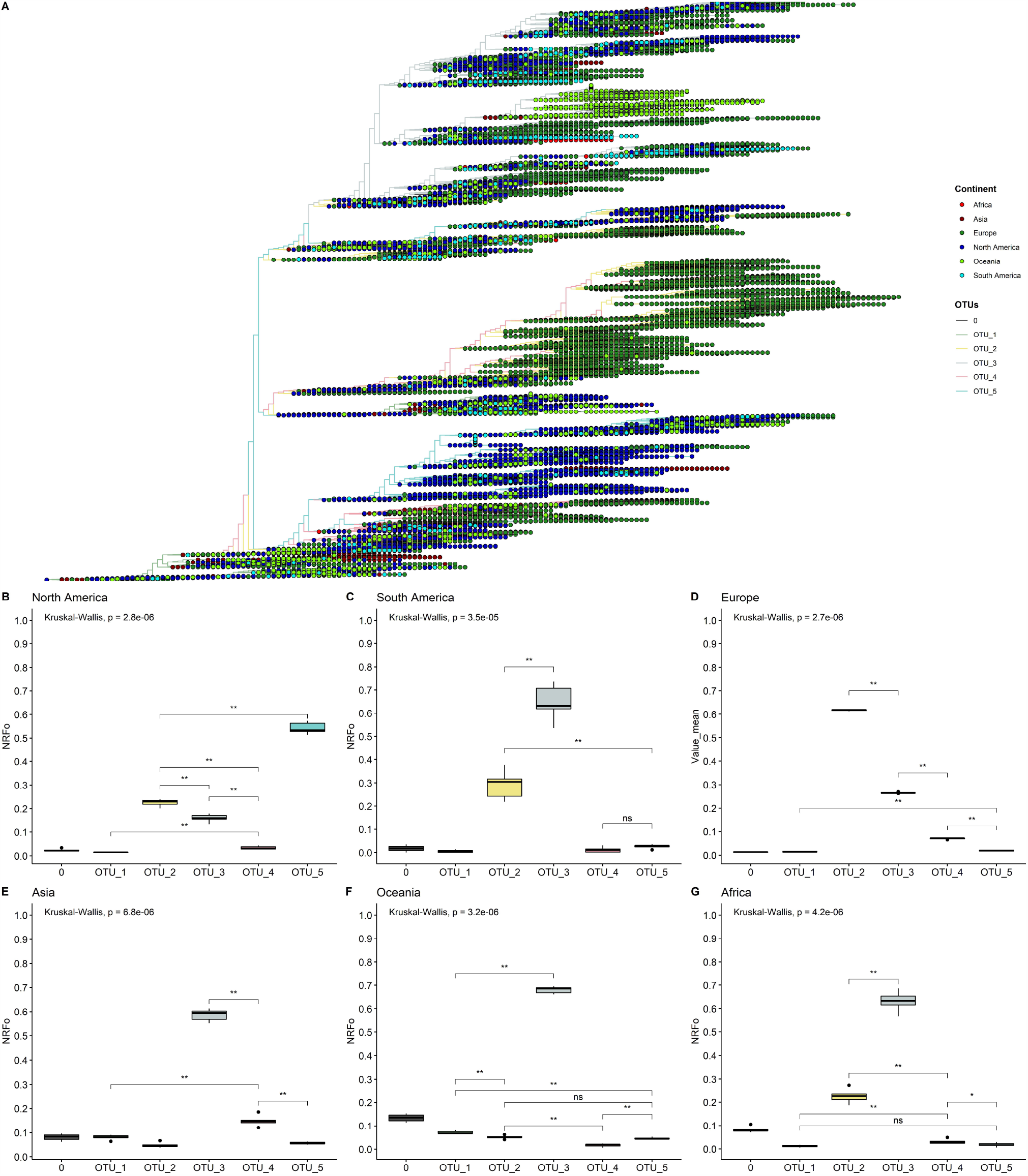
By cases normalized continent distribution of OTUs shows OTU_3 as the most prevalent in four of six continents. A) Unrooted tree of complete non-ambiguous genomes, tips were colored according to OTUs, and points in each tip were colored according to the continent. B-G) Boxplots of normalized relative frequencies of OTUs in each continent from December 2019 to November 2020 (B, North America; C, South America; D, Europe; E, Asia; F, Oceania; G, Africa).

To better analyze which were the most prevalent OTUs in each continent, we analyzed all the complete genomes in the GISAID database (171 461 genomes). In this analysis, we compared the mean of the frequency of OTUs normalized by cases in each continent of six randomly selected groups of genomes (see material and methods for more details).

This approach more clearly illustrates that OTU_5 was the most prevalent in North America, followed by OTU_2 and OTU_3, the least prevalent were OTU_1 and OTU_4 (Fig. 2.B). The first genomes in North America belonged to OTU_1 (Fig. S6). Since March, North America was dominated by OTU_5 (Fig. S6). OTU_5 has six of the nine high-frequency genomic variations described (all except those in N protein) (Fig. 1.A).

South America presents a greater OTU_3 frequency (Fig. 2.C) that was established in April (Fig. S5). This observation correlates well with studies focused in South America that detect the establishment of D614G mutation at the end of March (mutation presents in OTU_2, OTU_3, OTU_4 and OTU_5) and a high frequency of pangolin lineage B1.1 in Chile and in general in South America that contains the same characteristics mutations that our OTU_3 (Castillo et al. 2020, Franco-Muñoz et al. 2020). Unfortunately, few genomes are reported in South America for September, October, and November (24 genomes in total in the three months), hindering a correct analysis of frequencies in these months. Similarly, OTU_3 was most prevalent in Asia, Oceania, and Africa (Fig. 2.E, 2.F, and 2.G). With other OTUs with least than 0.3 NRFp (Fig. 2.E, 2.F, and 2.G). Wu et al. 2020 reports high incidence of mutations that define OTU_3 in Bangladesh, Oman, Russia, Australia and Latvia. At the haplotype level, OTU_3 presents mutations in the N protein that apparently increases the fitness of this group in comparison with OTU_2 (OTU_2 does not present mutations in N) (Fig. 1.A). Thus, four of the six continents analyzed presents an estimation of more than 50 % COVID-19 cases with a SARS-CoV-2 with the three mutations in the N protein. We, therefore, believe that is important to more deeply study if exists positive fitness implications for these mutations.

Europe presents an interesting pattern, it follows a similar pattern to South America, Asia, Oceania, and Africa until July (Fig. S6), with OTU_3 as the predominant. Then, in August, OTU_2 increased its frequency, and since September OTU_2 is the most prevalent in Europe. This could be caused by the appearance of mutations in the background of OTU_2 (such as those described in Justo et al. 2020) with greater fitness than those of OTU_3 or due to other effects (i.e., founder effects) after the relaxation of lockdown policies.

### Worldwide temporal distribution of OTUs

A rooted tree was estimated with the 109 953 genomes data set and labeled by date (Fig. 3.A). Here, we can observe that OTU_1 is mostly labeled with colors that correspond to the first months of the pandemic, expected due to its relation with the first genomes isolated. Clades, where OTU_2, OTU_3, OTU_4, and OTU_5 are the most prevalent, have similar distributions, with representatives mostly isolated since April. The biggest branch of OTU_2 presents a very specific temporal distribution with almost all the genomes isolated from September to November.

**Figure 3.**
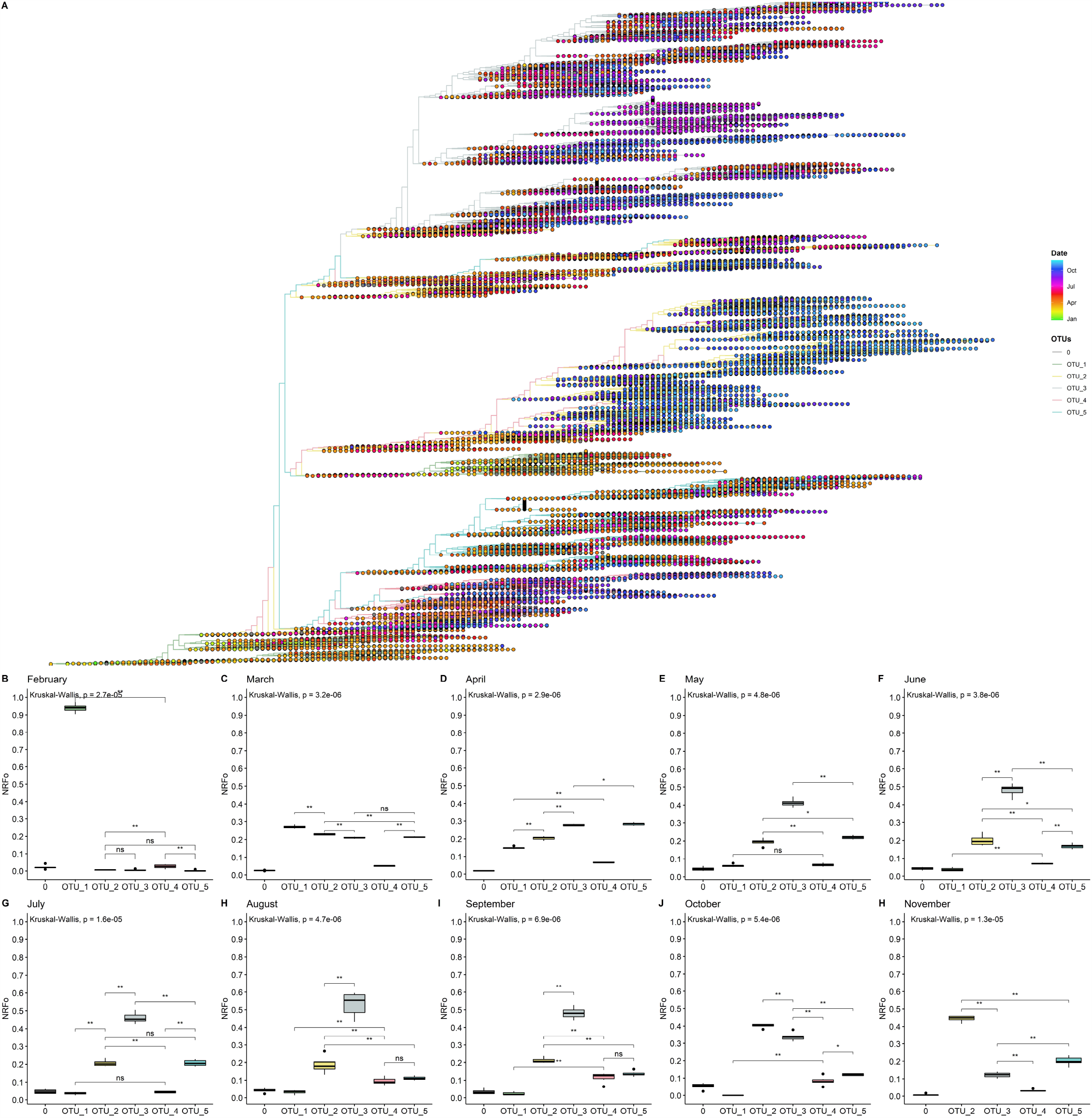
By cases normalized temporal distribution of OTUs showed OTU_3 as the most prevalent until September. A) Rooted tree of complete non-ambiguous genomes showing temporal distribution. Tips were colored by OTUs and points in each tip were colored according to the collection date. B-E) Boxplot of OTUs global distribution in each month (B, February; C, March; D, April; E, May; F, June; G, July; H, August; I, September; J, October; K, November).

To gain more insight into these patterns, we estimated the most prevalent OTUs in the world during each month of the pandemic following similar steps that those done for continents (see material and methods for details). In this analysis, we did not consider December and January that present all genomes except one belonging to OTU_1 and mainly from Asia (Fig.S6 and S7).

Analysis using the data of February from North America, Europe, and Asia showed that OTU_1 continued as the most prevalent in the world but with first isolations of OTU_2, OTU_3, OTU_4, and OTU_5 (Fig. 3.B). Analysis by continents showed that during this month Asia and North America still had higher proportions of OTU_1, but in Europe, a more homogeneous distribution of OTU_1, OTU_2 and OTU_3 was observed (Fig. S6).

In March, when the epicenter of the pandemic moved to Europe and North America, but cases were still appearing in Asia, OTU_2, OTU_3, and OTU_5 increased their prevalence but OTU_1 remained slightly as the most prevalent during this month (Fig. 3.C). Interestingly, OTU_4 remained in relatively low frequencies (Fig. 3.C). This month contains the more homogenous OTUs distribution in a worldwide context, but with some OTUs more prevalent in each continent (Fig. S6).

During April, OTU_1 continued its downward while OTU_3 and OTU_5 increased their presence (Fig. 3.D) probably due to its higher representation (compared to March) in several continents such as South America, North America, and Europe (Fig. S6). During this month, Africa showed a high prevalence of OTU_2 (Fig. S6). We also witnessed the establishment of OTU_3 in South America and OTU_5 in North America (Fig. S6).

May, June, and July showed a similar pattern, with OTU_3 as the most prevalent due to its high frequencies in South America, Oceania, and Europe (Fig. 3.E, 3.F, 3.G, and S6). North America maintains OTU_5 as the most prevalent and Oceania showed a relatively homogenous pattern. During these months, OTU_2 had intermediate frequencies in all continents resulting in intermediate frequencies all over the world (Fig. 3.E, 3.F, 3.G, and S6). OTU_1 and OTU_4 representatives were reported during these months but with very low frequencies.

In August and September, we detected a slightly higher frequency of OTU_4 compared to the previous months (Fig. 3H and 3I) with no significant differences with OTU_5. In September in Europe, OTU_3 stopped being the most frequent. Instead, OTU_2 was the most frequent in this month in Europe (Fig. S6). In October and November, OTU_2 has increased its frequency rapidly (Fig. 3.J and 3.H) mainly due to a large number of cases and reported genomes belonging to this OTU_2 in Europe in October and November. Due to the few genomes currently available in GISAID for all continents, except for Europe and North America during November, just these two continents were analyzed in the last month.

Also, it is important to mention that, there are not many enough genomes reported for September, October, and November for South America, so during these months OTUs frequencies of this continent were not considered.

### Age, Gender and Patient Status relation with OTUs

Relating the distribution of haplotypes according to patient information can help to determine the preference of some OTUs for some characteristics of the patients. Thus, we analyze OTUs distribution according to age, gender, and patient status information available as metadata in the GISAID database.

Unfortunately, just 26.11 % of the 171 461 genomes analyzed have age and gender information (Fig. S8). In the case of patient status information, we noted that GISAID categories are not well organized and we had to reclassify the information into three categories; Asymptomatic, Mild, and Severe (Fig. S9.A). Using this classification scheme, we found that 99.14 % (169 979 genomes) were not informative, 0.1 % (175 genomes) falls in the Asymptomatic category, 0.33 % (562 genomes) in the Mild category and 0.43 % (745 genomes) could be classified as Severe (Fig. S9.B).

Using this limited data, we attempt to determine whether any OTU causes an asymptomatic, mild, or severe infection more frequently. We look for significant differences between the relative frequencies of the OTUs in total samples and samples with known patient information. If we found differences, it would mean that some OTU could be more or less related to one type of infection. Here, we analyzed just the month-continent combination with at least 45 genomes with information of one type of infection and at least two times of genomes with any information (for example Asia – February has 58 Asymptomatic genomes and 613 total genomes). Ten combinations meet these criteria, one in the asymptomatic category, one in the mild, and eight in the severe. None of the OTUs frequencies in samples with patient status information were significative different from the frequencies in the total population of the month-continent analyzed (Fig. 4). Thus, we concluded that none of the OTUs are related to an asymptomatic, mild, or severe COVID-19, at least in the populations analyzed.

**Figure 4.**
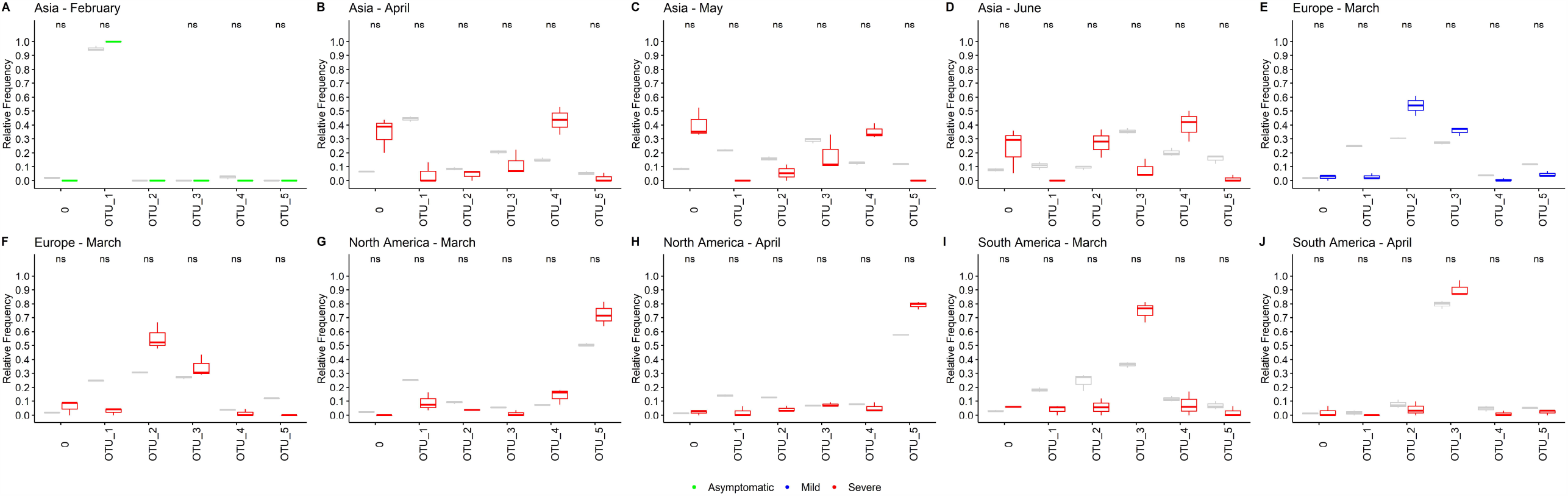
OTUs are not related to the COVID-19 severity. A-J) Ten different sample populations were analyzed, none of the OTUs frequencies shows significative differences between the total samples and samples taken from genomes with patient status information. Boxplots showed the distribution of three samples, total frequencies are showed in grey and frequencies from samples with patient status information are colored according the category (green, asymptomatic; blue, mild; red, severe).

Age information was also analyzed in the same manner. In general, although some differences were detected as significant, those were not consistently maintained between different populations analyzed (Fig. S10.A-J). Furthermore, none difference reaches a p-value less than (Except for OTU_4 in North America). Since heterogeneity between countries information is possible, we think that these small differences are more likely due to these heterogeneities and we cannot strongly conclude that some age groups are more related to a specific OTU. Additionally, a strong positive correlation between total relative frequencies of OTUs and relative frequencies by age groups in month-continent was found, meaning that those two frequencies are similar in most of the analyzed populations (Fig. S10.K)

A similar approach was done using gender information, but in this case, due to the greater quantity of information, we used more restrictive filter parameters. Thus, we selected country-month combinations with at least 250 genomes with male or female information and two times total genomes information (for instance USA – March has 2079 genomes from female patients and 9287 genomes with or without gender information). Again, we did not find OTU’s preference for a specific gender (Fig. S11).

### Description of the most frequent mutations

#### C241T

The C241T mutation is present in the 5’-UTR region. In coronaviruses, the 5’-UTR region is important for viral transcription (Madhugiri et al. 2014) and packaging (Masters. 2019). Computational analysis showed that this mutation could create a TAR DNA-binding protein 43 (TDP43) binding site (Mukherjee and Goswami. 2020), TDP43 is a well-characterized RNA-binding protein that recognizes UG-rich nucleic acids (Kuo et al. 2014) described to regulate splicing of pre-mRNA, mRNA stability and turnover, mRNA trafficking and can also function as a transcriptional repressor and protect mRNAs under conditions of stress (Lee et al. 2011). Experimental studies are necessary to confirm different binding constants of TDP43 for the two variants of 5’-UTR and its *in vivo* effects.

#### C1059T

Mutation C1059T lies on Nsp2. Nsp2 does not have a clearly defined function in SARS-CoV-2 since the deletion of Nsp2 from SARS-CoV has little effect on viral titers and so maybe dispensable for viral replication (Graham et al. 2005). However, Nsp2 from SARS-CoV can interact with prohibitin 1 and 2 (PBH1 and PBH2) (Cornillez-Ty et al. 2009), two proteins involved in several cellular functions including cell cycle progression (Wang et al. 1999), cell migration (Rajalingam et al. 2005), cellular differentiation (Sun et al. 2004), apoptosis (Fusaro et al. 2003), and mitochondrial biogenesis (Merkwirth and Langer. 2008).

#### C3037T

Mutation C3037T is a synonymous mutation in Nsp3; therefore, it is more difficult to associate this change with an evolutionary advantage for the virus. This mutation occurred in the third position of a codon. One possibility is that this changes the frequency of codon usage in humans increasing expression or any other of the related effects caused by synonymous codon change (some of them reviewed in Mauro and Chapel. 2014).

C3037T causes a codon change from TTC to TTT. TTT is more frequently present in the genome of SARS-CoV-2 and other related coronaviruses compared to TTC (Gu et al. 2014) but in humans, the codon usage of TTT and TTC are similar (Mauro and Chapel. 2014). The reason why TTT is more frequent in SARS-CoV-2 is unknown but seems to be a selection related to SARS-CoV-2 and not to the host. Another option is genetic drift.

#### C14408T

The C14408T mutation changes P323 to leucine in Nsp12, the RNA-dependent RNA polymerase of SARS-CoV2 (Fig. S12.A and B). P323 together with P322 ends helix 10 and generate a turn that is followed by a beta-sheet (Fig. S12.C). Leucine at position 323 could form hydrophobic interactions with the methyl group of L324 and the aromatic ring of F396 creating a more stable variant of Nsp12 (Fig. S12.E). In concordance with this, protein dynamics simulations showed a stability increase of the Nsp12 P323L variant (Chand and Azad. 2020). In the absence of P322, the mutation P323L would probably be disfavored due to the flexibilization of the turn at the end of helix 10. Experimental evidence is necessary to confirm these hypotheses and to evaluate their impact on protein function.

#### A23403G

An interesting protein to track is spike protein (Fig. S13.A) due to its importance in SARS-CoV-2 infectivity. It has been suggested that the D614G change in the S1 domain that results from the A23403G mutation generates a more infectious virus, less spike shedding, greater incorporation in pseudovirions (Zhang et al. 2020), and higher viral load (Korber et al. 2020).

How these effects occur at the structural level remains unclear, although some hypotheses have been put forward: 1) We think that there is no evidence for hydrogen-bond between D614 and T859 mentioned by Korber et al. 2020, distances between D614 and T859 are too long for a hydrogen bond (Fig S13.B), 2) distances between Q613 and T859 (Fig. S13.C) could be reduced by increased flexibility due to D614G substitution, forming a stabilizing hydrogen bond, 3) currently available structures do not show salt-bridges between D614 and R646 as proposed by Zhang et al. 2020 (Fig. S13.D).

#### G25563T

Orf3a (Fig. S14.A) is required for efficient in vitro and in vivo replication in SARS-CoV (Castaño-Rodriguez et al. 2018). It has been implicated in inflammasome activation (Siu et al. 2019), apoptosis (Chan et al. 2009), necrotic cell death (Yue et al. 2018) and has been observed in Golgi membranes (Padhan et al. 2007) where pH is slightly acidic (Griffiths and Simons. 1986). Kern et al. 2020 showed that Orf3a preferentially transports Ca+2 or K+ ions through a pore (Fig S14.B). Some constrictions were described in this pore, one of them formed by the side chain of Q57 (Fig. S14.C).

Mutation G25563T produces the Q57H variant of Orf3a (Fig. S14.C). It did not show significant differences in expression, stability, conductance, selectivity, or gating behavior (Kern et al. 2020). We modeled Q57H mutation and we did not observe differences in the radius of constriction (Fig. S14.C) formed by residue 57 but we observed slight differences in the electrostatic surface due to the ionizability of the histidine side chain (Fig. S14.D).

#### G28881A, G28882A, G28883C

N protein is formed by two domains and three disordered regions. The central disordered region named LKR was shown to interact directly with RNA (Chang et al. 2009) and other proteins (Luo et al. 2005), probably through positive side chains; also, this region contains phosphorylation sites able to modulate the oligomerization of N protein (Chang et al. 2013).

Mutation G28883C that changes a glycine for arginine at position 204 contributes one more positive charge to each N protein. Mutations G28881A and G28882A produce a change from arginine to lysine. These two positive amino acids probably have a low impact on the overall electrostatic distribution of N protein. However, change from R to K could alter the probability of phosphorylation in S202 or T205. Using the program NetPhosK (Blom et al. 2004), we observed different phosphorylation potential in S202 and T205 between G28881-G28882-G28883 (RG) and A28881-A28882-C28883 (KR) (Fig. S15). Other authors proposed that these mutations could change the molecular flexibility of N protein (Rahman et al. 2020).

## CONCLUDING REMARKS

Here, we present a complete geographical and temporal worldwide distribution of SARS-CoV-2 haplotypes from December 2019 to November 2020. We identified nine high-frequency mutations. These important variations (asserted mainly by their frequencies) need to be tracked during the pandemic.

Our haplotypes description showed to be phylogenetically consistent, allowing us to easily monitor the spatial and temporal changes of these mutations in a worldwide context. This was only possible due to the unprecedented worldwide efforts in the genome sequencing of SARS-CoV-2 and the public databases that rapidly share the information.

Our geographical and temporal analysis showed that OTU_3 is currently the more frequent haplotype circulating in four of six continents (Africa, Asia, Oceania, and South America), result that is in accordance with other studies (Mercatelli et al. 2020) that showed GISAID clade GR (that corresponds to our OTU_3) as the most prevalent in the world; however, they did not report the currently predominance of OTU_2 in Europe (clade G for GISAID). Intriguingly, OTU_3 never reached frequencies higher than OTU_5 in North America. In Europe, currently and different from the tendency from May to July, OTU_2 is now much more commonly isolated than OTU_3. Why mutations R203K and G204R have such frequencies in most of the continents, why in North America those mutations were not so successful and why currently Europe is dominated by OTU_2 are open questions. Some studies showed that at the moment there are not mutations that significative increase the fitness of the SARS-CoV-2 (Rasmussen et al. 2020, van Dorp et al. 2020).

Although OTU_1 was the only and the most abundant haplotype at the beginning of the pandemic, now its isolation is rare. This result shows an expected adaptation process of SARS-CoV-2. This enunciate does not mean that SARS-CoV-2 is now more infectious or more transmissible.

In the next months, these haplotypes description will need to be updated, identification of new haplotypes could be performed by combining the identification of new frequent mutations and phylogenetic inference. We will continue monitoring the emergence of mutations that exceed our proposed cut-off of 0.18 NRFp and this information will be rapidly shared with the scientific community through our web page (http://sarscov2haplofinder.urp.edu.pe/). This will also be accompanied by a continuous update of haplotypes information. During the peer-review process o this manuscript, we identify several other mutations near to the cut-off proposed that were reported in Justo et al. 2020.

Using information of specific populations we showed no preference for patient’s features (age, gender, or type of infection) by OTUs. Thus, mutations that define those haplotypes do not have a relevant impact on the severity of the disease neither are implied preferentially in infections to males, females, or age.

Finally, although more studies need to be performed to increase our knowledge of the biology of SARS-CoV-2, we were able to make hypotheses about the possible effects of the most frequent mutations identified. This will help in the development of new studies that will impact vaccine development, diagnostic test creation, among others.

## MATERIAL AND METHODS

### Normalized frequency analysis of each base or gap by genomic position

To perform the mutation frequency analysis, we first downloaded a total of 171 461 complete and high coverage genomes from the GISAID database (as of November 30^th^, 2020). This set of genomes was aligned using ViralMSA using default parameter settings, and EPI_ISL_402125 SARS-CoV-2 genome from nt 203 to nt 29674 as the reference sequence (Moshiri. 2020, Li. 2018). Subalignments corresponding to genomes divided by continent-month combinations was extracted and relative frequencies of each base or gap in each genomic position were calculated (*RF*_*p, m*−*c*_) using a python script. These relative frequencies were multiplied by the number of cases reported in the respective continent-month combination (*CN*_*m*−*c*_) obtaining an estimation of the number of cases that present a virus with a specific base or gap in a specific genomic position (*RF*_*p*_*CN*_*m*−*c*_). Finally, we added the *RF*_*p*_*CN*_*m*−*c*_ of each subalignment and divided it by the total number of cases in the world (∑_*m*−*c*_ *RF*_*0*_*CN*_*m*−*c*1_)/*TCN*_*w*_. This procedure allows us to obtain a relative frequency normalized by cases of each base or gap in each genomic position (*NRF*_*p*_). The number of cases of each country was obtained from the European Centre for Disease Prevention and Control: https://www.ecdc.europa.eu/en/publications-data/download-todays-data-geographic-distribution-covid-19-cases-worldwide. We used the number of cases of countries with at least one genome sequenced and deposited in GISAID database. Also, we just consider in the analysis month-continent combinations with at least 90 genomes sequenced.

### Phylogenetic tree construction

Using an alignment of the 109 953 complete, high coverage genomes without ambiguities, we estimated a maximum likelihood tree with Fasttree v2.1.10 with the next parameters: -nt -gtr - gamma -sprlength 1000 -spr 10 -refresh 0.8 -topm 1.5 close 0.75 (Price et al. 2009, Price et al. 2010), after the generation of the tree we improved topology using -boot 1000 and the first output tree as an input using -intree option. To generate the rooted tree (against EPI_ISL_402125) we used the R package treeio, and to generate tree figures with continent or date information by tip we used the ggtree package in R (Yu. 2020, Yu et al. 2017).

### OTUs determination

Mutations respect to EPI_ISL_402125 with NRF_p_ greater than 0.18 were extracted from the alignment of the non-ambiguous data set of 109 953 genomes and were associated with the whole-genome rooted tree using the MSA function from the ggtree package (Yu. 2020, Yu et al. 2017) in R. Then, we visually examined to identify the major haplotypes based in these positions, designated as OTUs (Operational Taxonomic Units). Haplotypes identification based in our NRFp calculation reduced the bias of the different number of genomes sequenced in each continent and each month by integrating the less biased information of the number of cases. Although, other biases are more difficult, if possible, to reduce or eliminate.

### Analysis of OTUs geographical distribution

In this analysis, we randomly separate the genomes into 6 samples of 28 576 genomes each. Genomes in each sample was divided by continents and by months. In these divisions, OTUs relative frequencies were calculated for each OTU in each month-continent combination (*O*_*n*_*F*_*m*−*c*_). Then, we multiplied these (*O*_*n*_*F*_*m*−*c*_) frequencies by the number of cases corresponding to the respective month-continent (*CN*_*m*−*c*_) to obtain an estimation of the number of cases caused by a specific OTU in a respective month-continent (*O*_*n*_*CN*_*m*−*c*_). After, these products were grouped by continents, and those from the same continent were added and then divided by the total number of cases in the continent analyzed (∑_*m*−*c*1_ *O*_*n*_*CN*_*m*−*c*1_)/*TCN*_*c*1_. Thus, obtaining a frequency normalized by cases for each OTU in each continent. Finally, following this procedure in each sample, we statistically compared the mean of those six samples using the package “ggpubr” in R with the non-parametric Kruskal-Wallis test, and pairwise statistical differences were calculated using non-parametric Wilcoxon test from the same R package. The number of cases of each country was obtained from the European Centre for Disease Prevention and Control: https://www.ecdc.europa.eu/en/publications-data/download-todays-data-geographic-distribution-covid-19-cases-worldwide. We used the number of cases of countries with at least one genome sequenced and deposited in GISAID database. Also, we just consider in the analysis month-continent combinations with at least 90 genomes sequenced.

### Analysis of OTUs temporal distribution

Following a similar procedure used in the geographical analysis, we now grouped the products *O*_*n*_*CN*_*m*−*c*_ by months, added them, and then divided by the total number of cases in the analyzed month (∑_*m*1−*c*_ *O*_*n*_*CN*_*m*1−*c*_)/*TCN*_*m*1_. As in the geographical analysis, the mean of the six samples was statistically compared using the same procedures and with exactly the same considerations of month-continent combinations.

### Analysis of age, gender, and patient status with OTUs distribution

We determine if OTUs have a preference for age or gender, or cause a COVID-19 with a specific severity. For patient status and age information we selected populations with at least 45 genomes in the category to analyze and at least two times the total number of genomes (for example Asia – February has 58 asymptomatic genomes and 613 total genomes). For the gender analysis, we selected sample populations with at least 250 genomes in the category to analyze and at least two times the total number of genomes (for example, USA – March has 2 079 genomes from female patients and 9287 genomes with or without gender information). In each selected sample we used the total data (all genomes corresponding to that continent-month combination) and the data with category information (for example male, female, asymptomatic, severe, 16-30 years, etc.). We randomly divided these two groups of genomes into three samples and calculated OTUs frequencies. The mean of the frequency of each OTUs was compared between the two groups using the non-parametric Wilcoxon or Kruskal-Wallis statistical test. In the case of age information, the relative frequencies of each OTUs of the total genomes and the genomes with category information were correlated using Spearman correlation. All plots were produced in R using “ggpubr” and ggplot2.

## DATA AVAILABILITY

The data that support the findings of this study comes from the GISAID initiative (Shu and McCaluey. 2017) (gisaid.org). Python and R scripts used in this study are available on request from the corresponding author upon reasonable request.

## Supporting information

Acknowledgements

## Acknowledgements

This manuscript has been released as a pre-print at https://doi.org/10.1101/2020.07.12.199414, (Justo et al.)

We are very grateful to the GISAID Initiative and all its data contributors, i.e. the Authors from the Originating laboratories responsible for obtaining the specimens and the Submitting laboratories where genetic sequence data were generated and shared via the GISAID Initiative, on which this research is based. Complete acknowledgements of the 171 461 genomes used are available in supplementary file (SF1-SF20).

We thank Professor Shaker Chuck Farah (Institute of Chemistry – University of Sao Paulo) for English writing corrections and helpful comments. Also, we thank Professors Aline Maria da Silva (Institute of Chemistry – University of Sao Paulo), Joao Renato Rebello Pinho (Albert Einstein Hospital – Sao Paulo) and PhD(c). Deyvid Amgarten (Albert Einstein Hospital – Sao Paulo) for its helpful comments. To the Ricardo Palma University High-Performance Computational Cluster (URPHPC) managers Gustavo Adolfo Abarca Valdiviezo and Roxana Paola Mier Hermoza at the Ricardo Palma Informatic Department (OFICIC) for their contribution in programs and remote use configuration of URPHPC. To Gladys Arevalo Chong for her figure style suggestions. To the Fundação de Amparo à Pesquisa do Estado de São Paulo (FAPESP) graduate scolarship (To S.J.A) 2015/13318-4 (to C. S. F.) and Universidad Ricardo Palma (URP) for APC financing.

## Supplemental Figures captions

**Table 1. Region of primers binding and amplification of nine diagnostic tests for SARS-CoV-2**.

**Table 2. Comparison between different nomenclatures of SARS-CoV-2 lineages**.

**Figure S1. Number of genomes sequenced by region is not correlated to the number of cases in the same region**. Each point in the plot represents a month-continent combination. There are continents with a high-number of cases but low number of sequenced genomes and inversely, there are continents with relatively few cases but with a large number of sequenced genomes.

**Figure S2. Normalized Relative Frequency of each nucleotide by position (NRF**_**p**_). The frequency of each nucleotide in each position was normalized by the number of cases in each continent-month pairs to reduce the bias produced by the different number of sequenced genomes in different months and different continents. In A, Labels are showed for NRF_p_ greater than 0.18. B and C showed different scales of positions with less than 0.18 NRF_p_.

**Figure S3. Temporal distribution by day, continent, and OTUs**. Each point in the plot represents one of the 171 461 SARS-CoV-2 genomes analyzed. Points are colored depending on the OTU. Y-axis divides the points in continent and each column represents a day from December 16 to July 23.

**Figure S4. Global NRFp of the nine most frequent mutations by month**. Mutations that define OTU_2 (C241T, C3037T, C14408T, A23403G) showed very similar frequencies indicating that genomes with three, two or one of these mutations are rare. The same for mutations that define OTU_3 (G28881A, G28882A, G28883C). Mutations that define OTU_4 (C1059T) and OTU_5 (G25563T) have similar but not identical distributions.

**Figure S5. Month and continent distribution of the 171 461 SARS-CoV-2 genomes analyzed**. A) Bars represent genome count in each continent analyzed. Europe and North America are overrepresented in the database. B) Bars in B represent genomes count by month. March, April and October are the best represented months. Bars are labeled by percentage and below by the exact counts.

**Figure S6. Temporal distribution by month, continent, and OTU**. Each point in the plot represents a genome and is colored depending on OTU. Points are grouped by continent (Y-axis) and month (x-axis). We saw how haplotypes populations changes during time; for example, OTU_1 seems the most common during the first months (December, January, and February).

**Figure S7. Distribution of OTUs in January**. Bar plot of a count of complete genomes isolated in January and deposited in the GISAID database. Most of these genomes belonging to OTU_1, a small fraction corresponds to unclassified genomes and one to OTU_2

**Figure S8. Approximately 74 % of the genomes in GISAID database does not have gender information**. The plot shows gender distribution of the 171 461 SARS-CoV-2 genomes analyzed. Bars represent genomes count in Male, Female or unknown categories.

**Figure S9. More than 90 % of the genomes in the GISAID database does not have an informative description of patient status**. A) Table showing which GISAID categories were recategorized in the Asymptomatic, mild or severe categories. All the other genomes were classified as non-informative. B) Distribution of 171 461 genomes in patient status categories (Asymptomatic, Mild, Severe or No informative).

**Figure S10. Age groups are not robustly related to OTUs**. A-J) Ten populations were selected to analyze if OTUs frequencies in an age group is significative different to OTUs frequencies in the total population. None OTU showed a repetitive preference for an age group in the populations analyzed, boxplots are colored by age groups, all means frequencies in the total population (ns, p>0.05; *, 0.05>p>0.01; **, 0.01>p>0.005; ?, not analyzed). K) Correlation between relative frequencies of OTUs in a specific age group with OTUs frequencies in the whole population. Spearman correlation showed an R value of 0.94 meaning a positive correlation that supports the conclusion that no significative differences exist between OTUs frequencies in age groups compared to the whole population.

**Figure S11. OTUs do not have preference for males or females**. A-K) Boxplots of OTUs frequencies from female populations compared to OTUs frequencies in the whole population. None significant difference was observed. L-V) The same as A to K but whole population compared to male populations. Again, no significant differences were observed. Concluding that OTUs do not show gender preferences.

**Figure S12. P323L could impact the stability of Nsp12 without disturbing its overall structure**. A) Structure of RNA-dependent RNA polymerase complex (PDB ID: 6YYT). Chains (Nsp12, Nsp7, Nsp8, RNA) are distinguished by colors. Helix 10, Beta-sheet 3, Turn 10-3, and P323 also are differentially colored. B) Structure in A rotated 90 degrees. C) Zoom of the red box in B showed P322 and P323 in the center of Turn 10-3. D) Turn 10-3 with side chains of P323, L324, and F396 in sphere representation to highlight the distance between side chains of P323 and L324. E) P323 in D was computationally replaced by L323. Now, distances between the methyl group of leucine are shorter with L323.

**Figure S13. Structural hypotheses about D614G mutation in Spike protein**. A) Structure of the open state of Spike trimer (PDB ID: 6YVB) colored by domains. B) Distances between side chains of two possible rotamers of D614 (1’-D614 and 2’-D614) and T859. Except for 1’-D614 and carbonyl group of T859, the other distances seems to be large to form a hydrogen bond.

C) Distances between side chains Q613 and T859. These distances are also large to form hydrogen bonds. D) R646 points to the opposite side of D614 showing that there is no salt bridge. B, C, and D show electron density maps of the side chains of the labeled residues.

**Figure S14. Orf3a Q57H does not modify pore constriction distances but electrostatics distribution**. A) Structure of the Orf3a dimer (PDB ID: 6XDC) colored by domains. The right of A shows the same structure but in an upper view. B) Orf3a showing the central pore, in the red box the section corresponding to the fifth pore constriction. C) zoom of the red box in B, above we showed Q and H variants superposed. Below we show a transversal cut of the pore near to the fifth. The pore radius in two variants is similar. D) Electrostatic surface maps of Q57 and H57 variants in two different pHs (7 and 6). Residues Q57 and H57 are shown in stick representations to point the fifth constriction. We show a slightly more positive region at the height of the fifth constriction.

**Figure S15. Mutants in R203 and G204 of Nucleocapsid generate differences in Phosphorylation potential on S202 and T205**. Bar plot showing the phosphorylation potential calculated in NetPhosK for the 4 possible nucleocapsid variants. We can see that phosphorylation potential by PKC is lower for RG than for KR in S202. On the other hand, T205 has greater phosphorylation potential by an unspecific kinase (unsp) in RG than in KR. Phosphorylation in S202 and T205 by unsp or PKC respectively is apparently not affected by these mutations.

